# A Trait Syndrome Ties Cell Morphology to Glycolysis Across the Yeast Subphylum

**DOI:** 10.1101/2025.10.27.684252

**Authors:** Linda C. Horianopoulos, Christina M. Chavez, Antonis Rokas, Chris Todd Hittinger

## Abstract

Traits that co-vary across species can provide fundamental insights into the trade-offs and constraints that govern their evolution. It was recently reported that glucose uptake rates (GUR) are inversely correlated with the cell surface area-to-volume (SA:V) ratio across 11 yeast species. Here we substantially expand this analysis to 282 species to test whether the GUR-SA:V correlation generalizes across the ancient Saccharomycotina yeast subphylum and to determine the contribution of shared evolutionary history to the co-variation of these two traits. Using regression models that account for co-variation due to phylogeny, we found that extracellular acidification rates (ECAR, which we used as a scalable proxy for GUR) had a weak, but significant, correlation with SA:V across Saccharomycotina. We found additional weak, but significant, correlations between ECAR with genome sizes and growth rates. Our findings largely agree with the recently reported correlations between GUR and SA:V ratio, but they also show that there are likely several other associated variables, including genome size. Specifically, yeasts that consume glucose faster tend to have lower SA:V, faster growth rates, and larger genomes. Our results suggest that a trait syndrome governs several metabolic, genomic, and morphological traits across the yeast subphylum.

## Introduction

The Saccharomycotina yeast subphylum includes over a thousand species and displays impressive genetic and phenotypic diversity^1–5^. The ability of most yeast species to grow well in axenic culture enables both the characterization of many traits and the exploration of biological principles that govern their variation^2^. Such studies have revealed that certain traits are consistently associated with each other and may collectively contribute to trait syndromes^6^. These trait syndromes can be driven by intrinsic factors (e.g. genomic characteristics) or extrinsic factors (e.g. varying resource availability)^2,7^.

Recently, Li et al.^8^ reported the interesting finding that glucose uptake rate (GUR) is inversely correlated with surface area-to-volume ratio (SA:V) across 16 yeast strains in 10 Ascomycota species (subphyla Saccharomycotina and Taphrinomycotina) and one Basidiomycota species (subphylum Pucciniomycotina). Parallel to and independently from Li et al.^8^, we had measured glucose-induced extracellular acidification rates (ECAR) as a proxy for glycolytic rates^9^ and characterized yeast phenotypic variation by compiling data on asexual budding cell size and shape^3^ of 282 diverse yeast species within Saccharomycotina. Our dataset is uniquely suited to test whether the correlation reported by Li et al.^8^ holds true across an entire yeast subphylum while controlling for phylogenetic co-variance. We also considered two additional traits, genome size and rates of growth on glucose, which were previously collected for these yeast species as part of the Y1000+ Project (http://y1000plus.org)^2^.

## Results and Discussion

First, we calculated the surface area, volume, and SA:V of each yeast species and retrieved their ECAR, genome size, and growth rate values^2,9^ (Table S1, see Methods). We then calculated the phylogenetic signal of each trait using Pagel’s lambda and found that all traits had high and significant phylogenetic signal values (ECAR: λ = 0.815, p-value = 1.14E^-49^; SA:V ratio: λ = 0.846, p-value = 1.23E^-19^; genome size: λ = 0.975, p-value = 1.07E^-42^; and growth rate on glucose: λ = 0.725, p-value = 5.78E^-7^). The high phylogenetic signal values suggest that some of the variation in each trait can be attributed to shared evolutionary history. Thus, we applied phylogenetically corrected regression models (see Methods) to determine the relationships that exist between ECAR and SA:V and genome size (n = 282 species), as well as between ECAR and growth rate on glucose (n = 231 species). We found that ECAR was positively correlated with SA:V (phylogenetic signal λ of the model = 0.808, R = 0.23, p-value = 9.93e-5, Figure 1A). Since ECAR is negative (faster glucose-induced acidification) when GUR is high, these results support the main finding of Li et al.^8^ and reveal that the inverse correlation between GUR and SA:V holds across a much larger group of yeast species and that it cannot be explained solely through phylogenetic co-variance.

**Figure 1.**
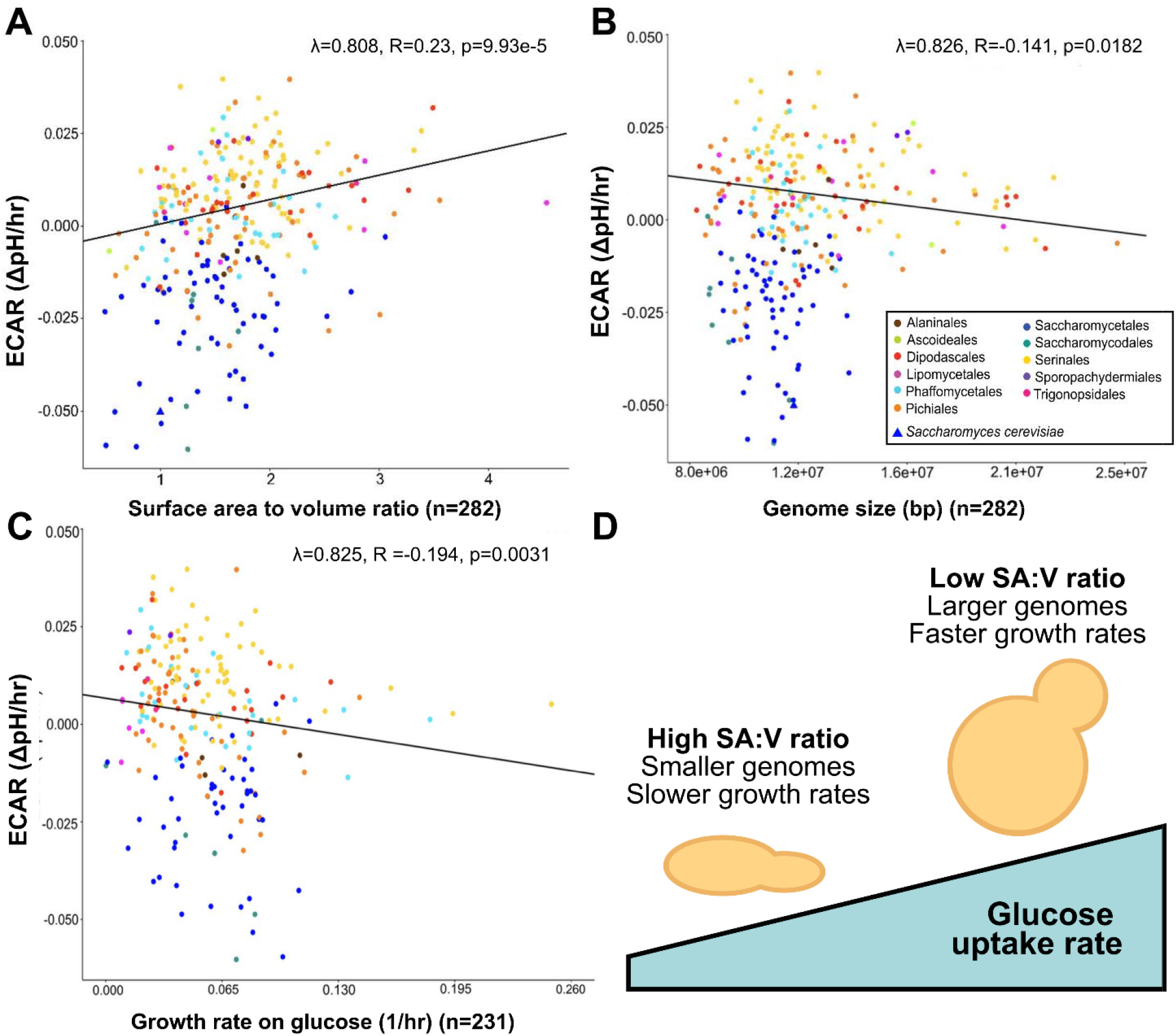
Glycolytic rate is significantly associated with cell morphology, genome size, and growth across the Saccharomycotina yeasts. (A-C) Scatter plots show significant correlations between ECAR (extracellular acidification rates, measured as a proxy for glycolytic rates) and: (A) cell surface area-to-volume ratio; (B) genome size; and (C) growth rate on glucose. Each dot represents a single yeast species colored according to its taxonomic order as labeled (B). *S. cerevisiae* is represented by a triangle in panels (A) and (B). The R and p values are based on a phylogenetic generalized least squares model, which accounts for correlation stemming from shared phylogenetic history. (D) A schematic model proposes a trait syndrome associated with a rapid glucose uptake rate among diverse yeast species.

Typically, smaller cells with larger SA:V ratios have been found to exhibit higher glycolytic rates due to increased transport capacity at the cell surface^10,11^. Smaller SA:V ratios may also place fewer resource demands on cell wall and membrane biosynthesis. However, an alternative model suggests that efficient scaling of biosynthesis and sufficient protein production are key to cell enlargement and rapid metabolism^12^. Our findings are largely consistent with this second model as cells with lower SA:V ratios generally had higher glycolytic rates. We speculate that, in yeasts, larger cells have more space in the cytosol for the catabolism of glucose, thereby enabling a stronger pull through glycolysis and consequently rapid GUR and growth rates. This glycolytic effect may be particularly applicable to Crabtree/Warburg-positive yeasts, which ferment in the presence of oxygen and have been proposed as an exception to the classic view that high SA:V is necessary for high GUR^10^. Overall, this model proposes that the glycolytic rate of yeasts growing on glucose is more influenced by enhanced glycolytic flux enabled by increased cytosolic volume than by nutrient import.

In contrast to the findings of Li et al.^8^, our larger dataset also showed significant negative correlations between ECAR and genome size (λ = 0.826, R = -0.141, p-value = 0.0182, Figure 1B) and between ECAR and growth rate on glucose (λ = 0.825, R = -0.194, p-value = 0.0031, Figure 1C). The contrast between our findings and those of Li et al.^8^ is likely explained by the enhanced statistical power of our dataset. Previous work showed that a larger genome size was associated with a generalist phenotype in Saccharomycotina yeasts, wherein generalists grew on a wider range of carbon sources and had faster growth rates^2^. Our data further show that yeast species with larger genomes have increased glucose uptake rates and faster rates of growth on glucose. Therefore, yeasts with larger genomes, such as those with whole genome duplications^13^, could be promising for industrial applications requiring rapid metabolism.

Although we interpret our correlations as a relationship between GUR and cell SA:V ratio, these results must be interpreted with caution since metabolic differences, particularly in acid production, may influence the validity of using ECAR as a proxy for GUR across different species. To supplement our finding of an association between low rates of ECAR and increased SA:V ratio, we considered the recently reported discovery of a group of yeasts within the genus *Saturnispora* that evolved elevated glycolytic rates^9^. Specifically, we noticed that two congeneric species, *Saturnispora silvae* and *Saturnispora dispora*, displayed both contrasting ECAR and GUR phenotypes and contrasting cellular morphologies (Figure 2A), which are consistent with the observations of Li et al.^8^ and our observations above. Moreover, a *Sat. dispora* deletion mutant lacking the gene encoding the transcription factor Gal4 had both decreased glucose consumption rate and decreased ECAR (despite increased acetate accumulation)^9^. We also found that this deletion mutant had an increased SA:V ratio when compared to the wild-type control (p = 0.0011, Figure 2B, Table S2), which is again consistent with the trend observed between metabolism and morphology in yeasts. These experimental observations show that the correlations observed above have predictive value in species beyond those studied by Li et al.^8^

**Figure 2.**
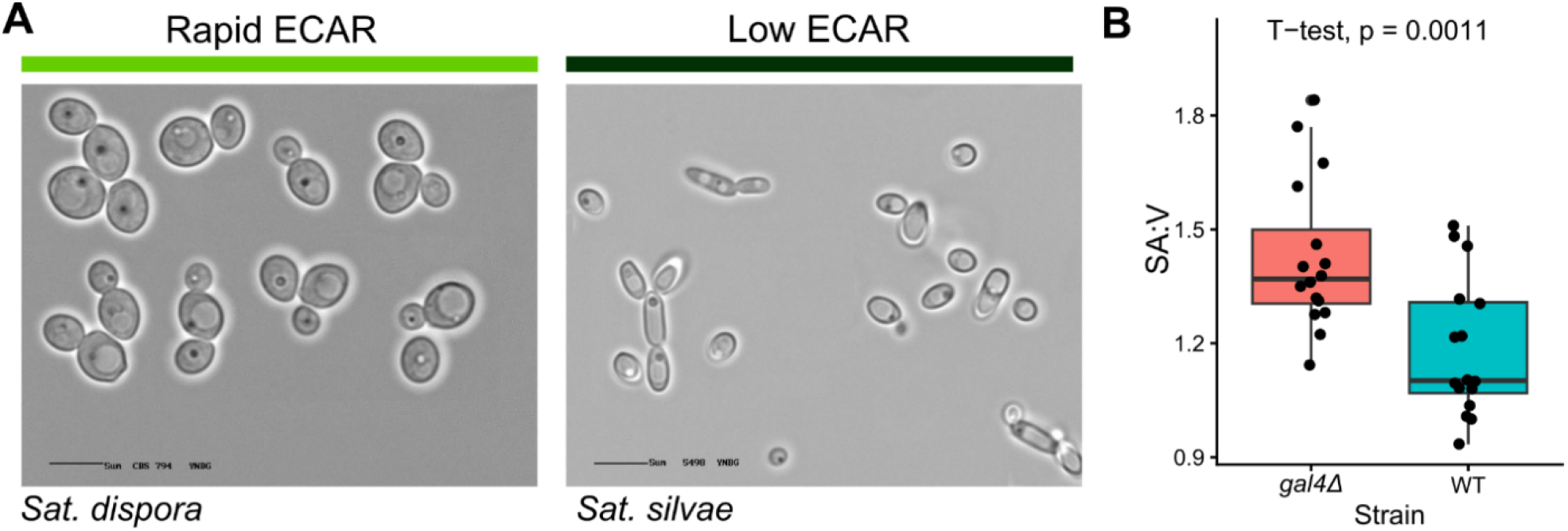
Morphology changes across varying glycolytic rates and a *GAL4* mutant in *Saturnispora dispora*. (A) Microscopy images (from https://theyeasts.org) of rapid- (*Sat. dispora*) and low-ECAR (*Sat. silvae*) yeasts reveal clear morphological differences that are consistent with the findings that glucose uptake rate is correlated with cell morphology. (B) A *Sat. dispora* mutant lacking *GAL4* has decreased glucose consumption^9^ and displays a higher surface area-to-volume ratio compared to the control strain. For the boxplots, center lines represent the median, bounds of the boxes represent interquartile ranges, whiskers represent the spread of the data, and each dot represents the measurement of one cell (n=16). The p value is the result of a two-sided t-test.

## Conclusions

Altogether our results and previous work lead us to propose a trait syndrome in which glycolytic rate is associated with multiple morphological and genomic features across the yeast subphylum. We found significant, but weak, correlations across several traits of interest, which suggests the existence of a complex trait syndrome, albeit one with considerable unexplained variance. Further, the associations between metabolism and genome size could suggest that gene acquisitions, gene family expansions, or genome duplications may be, at least in part, influencing glucose uptake rates and cell size as these processes are known to have impacts on metabolism^13–17^. Finally, our results raise important questions with respect to the mechanisms that link primary carbon metabolism to the regulation of cell morphology, whether genomic features other than size contribute to the relationship between these traits, and the environments that select for this trait syndrome. The findings reported here and by Li et al.^8^, alongside the rich genomic and phenotypic dataset available for the Saccharomycotina subphylum^2^, are poised to guide experiments that will uncover these mechanisms.

## Methods

The raw data for each trait in each species were collected and compiled from previously published work: ECAR^9^, morphology^3,5^, growth rate on glucose, and genome size^2^. The methodology for measuring ECAR is described in detail elsewhere^9^, but we would like to briefly highlight some key points considered to improve the reliability of this measurement as a proxy for glycolytic rate. All yeast species pre-cultures were washed and normalized by OD_600nm_ in water to remove any accumulated extracellular metabolites. Additionally, all measurements were taken in the first 90 minutes after glucose addition so that extracellular metabolites had not yet accumulated. During these early timepoints, acidification is primarily due to proton extrusion upon the uptake and cytosolic phosphorylation of glucose to buffer cytosol acidification^18^. The surface area and volume were calculated based on a spherical or ellipsoid categorical assignment of each yeast^5^.

To investigate the role of shared evolutionary history in the variation of each trait, we calculated Pagel’s λ using the phylosig() function of the *R* package phytools version 2.4-4 with a set seed^19^. Since each of these traits each had a high phylogenetic signal value, we applied phylogenetically corrected linear regression models to determine the relationships that exist between traits using the *R* package phylolm version 2.6.5 function with model set to ‘lambda’^20^.

The SA:V ratio of *Sat. dispora* strains was measured from images of cells captured with an EVOS FL microscope with 100x oil immersion plan apochromat objective lens. The length and width of cells were measured using Fiji^21^, and the data were processed using *R* v. 4.3.2 and visualized using ggplot2 v3.5.2^22^

## Acknowledgements

We would like to thank members of the Y1000+ Project (http://y1000plus.org) for helpful discussions. This project was supported by the National Science Foundation under Grants No.

DEB-2110403 (to C.T.H.); DEB-2110404 (to A.R.); in part by the Great Lakes Bioenergy Research Center, U.S. Department of Energy, Office of Science, Biological and Environmental Research Program under Award Number DE-SC0018409 (of which C.T.H. is a co-investigator); the National Institute of Food and Agriculture, United States Department of Agriculture, Hatch project 7005101 (to C.T.H.); and the Vilas Trust Estate. Research in A.R.’s lab is also supported by the National Institutes of Health/National Institute of Allergy and Infectious Diseases (R01 AI153356). L.C.H. was supported by a Natural Sciences and Engineering Research Council of Canada (NSERC) postdoctoral fellowship.

## Author contributions

L.C.H, C.M.C, A.R., and C.T.H. conceptualized this project. L.C.H and C.M.C. developed the methodology, curated and analyzed the data, and prepared the figures under the supervision and guidance of A.R. and C.T.H. L.C.H and C.M.C wrote the manuscript, and A.R. and C.T.H. reviewed, edited, and approved the manuscript. L.C.H., A.R., and C.T.H. acquired funding for this project.

## Declaration of Interests

A.R. is a scientific consultant of LifeMine Therapeutics, Inc. All other authors declare no competing interests.

## Notes

### Summary of Updates

We used different software to correct PGLS R values. We added a second figure containing new confirmatory data.

